# Increased numbers of CD4^+^ T-cells in the hypocretin/orexin region of Narcolepsy Type 1

**DOI:** 10.1101/2025.08.08.669086

**Authors:** Ling Shan, Milo Fonville, Mabel Hoekstra, Joost Smolders, Dick F. Swaab

**Affiliations:** Neuropsychiatric Disorders Group, Netherlands Institute for Neuroscience, an Institute of the Royal Netherlands Academy of Arts and Sciences, 1105 BA Amsterdam, the Netherlands; Neuroimmunology Research Group, Netherlands Institute for Neuroscience, an Institute of the Royal Netherlands Academy of Arts and Sciences, 1105 BA Amsterdam, the Netherlands; Departments of Neurology and Immunology, Erasmus MC, University Medical Center Rotterdam, 3015 CN Rotterdam, the Netherlands

**Keywords:** narcolepsy type 1 (with cataplexy NT1), CD4^+^ T-cells, neuropathology, hypocretin/orexin, hypothalamus

## Abstract

Narcolepsy type 1 (NT1) is proposed to be an autoimmune disorder targeting hypothalamic hypocretin (orexin, Hcrt) neurons. Hcrt reactive T-cells have been identified in blood and cerebrospinal fluid (CSF) of NT1 patients. However, it remains unknown whether T-cells infiltrate the brain. Since T-cells can be retained for a lifetime in tissues as tissue resident memory T-cells after primary antigen exposure, we now systematically assess the presence of CD4^+^ and CD8^+^ T-cells in NT1 brains to determine their regional distribution and potential autoimmune involvement in NT1 neuropathology.

We immunohistochemically stained and quantified CD4^+^ and CD8^+^ T-cells in post-mortem brain tissue of NT1 patients (n=5) and matched controls (n=5) in the Hcrt region, paraventricular nucleus (PVN) and median eminence (ME) as well as in the substantia nigra (SN) and locus coeruleus (LC). To phenotypically characterize CD4^+^ T-cells, we performed double staining with CD49a or C-X-C chemokine receptor type 6 (CXCR6). Furthermore, we stained for fibrinogen to estimate blood-brain barrier integrity, as well as microglia markers and an astrocyte marker to evaluate acute immune reactivity in the Hcrt region of NT1 brains.

In NT1 there was an 11-fold increase in total number of CD4^+^ T-cells in the Hcrt region, but not in the PVN, ME, nor SN or LC. These CD4^+^ T-cells exhibited tissue residential memory features with double staining with CD49a or CXCR6. There were no changes in blood-brain barrier integrity, microglia and astrocyte staining intensities between NT1 and controls. In addition, the total number of CD4^+^ T-cells in the Hcrt region showed significant negative correlations with mean sleep latency in NT1 cases.

Our findings suggest an enrichment of CD4^+^ T-cells specifically in the Hcrt region of NT1 indicating prior local antigen engagement. Moreover, greater CD4^+^ T-cell presence in the Hcrt region of NT1 was associated with increased symptom severity, as reflected by shorter sleep latency. These data support the hypothesis that CD4^+^ T-cells infiltrate the Hcrt region, where they may contribute to the autoimmune process that initiates NT1.

## Introduction

Narcolepsy type 1 (narcolepsy with cataplexy, NT1) is a rare, disabling neurological disorder characterised by impairment of sleep-wake functions, including excessive daytime sleepiness, cataplexy (sudden loss of muscle tone triggered by emotions), sleep paralysis and disturbed nocturnal sleep (Bassetti *et al*., 2019). The disease is mainly caused by a lack of hypocretin (orexin, Hcrt) neurons in the hypothalamus (Peyron *et al*., 2000; Thannickal *et al*., 2000; Mignot *et al*., 2002), while recently it has been shown that NT1 is also associated with the loss of corticotrophin-releasing hormone (CRH) neurons in the paraventricular nucleus (PVN) (Shan *et al*., 2022).

An auto-immune mediated etiology has been long hypothesized based mainly on human leukocyte antigen (HLA) subtype DQB1*06:02, T-cell receptor genetics (JUJI *et al*., 1988; Mignot *et al*., 2001; Hallmayer *et al*., 2009; Han *et al*., 2014; Ollila *et al*., 2023) and epidemiological risks such as winter upper airway infections, H1N1 influenza infections and certain vaccines (Han *et al*., 2011; Stowe *et al*., 2020; Ollila *et al*., 2023). However, direct human brain evidence to support this presumed autoimmune attack causing NT1 is lacking. Microglia marker (Allograft inflammatory factor 1 also known as IBA-1) is unchanged in the Hcrt region (Honda *et al*., 2009) or PVN (Shan *et al*., 2022) of NT1 postmortem human brains compared to matched controls. Moreover, glial fibrillary acidic protein (GFAP), an astrocyte marker, showed inconsistent results in the NT1. It has been reported as unchanged (Honda *et al*., 2009) in NT1 or significantly increased (Thannickal *et al*., 2003). The inconsistent postmortem human brain observations could be caused by a long disease duration. However, a positron emission tomography (PET) study, on 41 NT1 cases with recent disease onset showed no difference in microglia density in hypothalamus as well as in other brain regions (Barateau *et al*., 2024). It may be pointed out that other innate immune factors may be involved in disease pathophysiology. In particular, blood or cerebral spinal fluid CD4^+^ (Latorre *et al*., 2018; Jiang *et al*., 2019) and CD8^+^ (Pedersen *et al*., 2019) T-cells from NT1 patients were shown to autoreact with Hcrt peptides. However, the question remained to be answered whether T-cells entering NT1 patient brains play a direct role in the autoimmune processes or peripheral T-cells interacted with Hcrt peptides that are released into the circulation following the loss of Hcrt neurons as has been shown for other neuronal proteins (Van Zwam *et al*., 2009).

The human central nervous system is populated by CD8^+^ and CD4^+^ T-cells with a tissue-resident memory T-cell phenotype (Smolders *et al*., 2018; Hsiao *et al*., 2023, 2025). In other tissues, such as the lung, gut, and skin, tissue resident memory T-cell populations arise at sites of initial antigenic challenge and are maintained lifelong at these sites (Mueller and Mackay, 2016). Given the strong association between NT1 and the HLA-DQB1*06:02 allele, we hypothesized an enrichment of CD4^+^ T-cells in NT1-relevant regions of the hypothalamus which reflect prior engagement with local Hcrt-neuronal antigens in NT1. The primary goal of the current neuropathological study was to characterize the density of CD4^+^ and CD8^+^ T-cells within the Hcrt neuron region of the hypothalamus in NT1 patients to explore their potential direct role in Hcrt neuron loss. To answer whether T-cell enrichment is specific to the Hcrt region, we also systematically quantified the number of T-cells in the adjacent PVN and median eminence (ME) regions, where we recently reported a loss of CRH neurons and fibers (Shan *et al*., 2022), as well as in the substantia nigra (SN) and locus coeruleus (LC). In addition, the CD8/CD4 ratio in hypothalamus, SN and LC were investigated to test whether HLA-DQB1*06:02 carriership in NT1 patients alone influenced the ratio compared to non-carrier controls. Furthermore, to assess blood-brain barrier integrity we stained for fibrinogen, while to evaluate acute immune responses we used IBA-1 and Human Leukocyte Antigen-DR (HLA-DR) as microglia markers, as well as Glial Fibrillary Acidic Protein (GFAP) as the astrocyte marker and the B-cell marker CD79a. Since current results in NT1 showed increased CD4^+^ T-cell in the Hcrt region, we also tried a phenotypic characterization of those CD4^+^ T cells to double stain with CD49a (also known as Integrin alpha-1) or C-X-C chemokine receptor type 6 (CXCR6).

## Materials and Methods

All the postmortem human brain tissue, clinical diagnosis and neuropathological reports were obtained from the Netherlands Brain Bank (NBB). The donor or the next of kin provided permission to the NBB to use the brain tissue, clinical and neuropathological information for scientific research. The NBB procedures concerning “Donation of brain material for scientific research” were approved by the independent Review Board (IRB) of the Vrije University Medical Center under reference number 2009/148. Moreover, all operations of NBB was in compliance with European ethical and legal guidelines (Klioueva *et al*., 2015, 2018). Detailed information on the brain samples and clinical-pathological information was published in our previous study (Table 1 and Supplementary Table 1) (Shan *et al*., 2023). In short, four hypothalamic from patients with NT1 with positive HLA-DQB1*06:02, strong diminished Hcrt and CRH cells have been reported previously (Shan *et al*., 2022). In addition, one HLA-DQB1*06:02 positive NT1 patient (NBB2010-064) was first diagnosed as an NT1 patient with cataplexy at age of 61, but after eight year of chronic treatment with opiates as pain relief, the patient was reclassified as having idiopathic hypersomnia without cataplexy (Thannickal *et al*., 2018). This case has relatively more hypocretin neurons (33% of control total number of Hcrt neurons) than regular NT1 cases (3% of control) (Shan *et al*., 2022). We presumed that the residential memory T-cell numbers in this specific case might shed light on the relationship between T-cells and Hcrt neurons. Five non-neurological controls matched for sex, age, postmortem delay, brain weight and circadian clock time of death, cerebral spinal fluid (CSF)-pH, Braak stage for tangles (Alzheimer’s, AD stage; (Braak *et al*., 1996)), amyloid stage, Braak Lewy body stage (Parkinson’s, PD stage;(Braak *et al*., 1998)) as well as fixation time.

**Table 1.** Clinical-pathological information.

| NBB | Age | Sex | PMD | pH | BW | B-AD | A | B-PD | Cause of Death | Relevant Medication |
| --- | --- | --- | --- | --- | --- | --- | --- | --- | --- | --- |
| **Narcolepsy type 1** |  |  |  |  |  |  |  |  |  |  |
| 2008-023 | 66 | f | 420 | 6.6 | 1175 | 1 | A | 0 | Heart failure | Amphetamine;<br>Modafinil; Sodium oxybate |
| 2018-018 | 82 | f | 330 | 6.5 | 1165 | 2 | O | 0 | Pneumonia | Modafinil |
| 2018-091 | 71 | f | 395 | 6.3 | 1215 | 4 | A | 0 | Heart failure, renal insufficiency | Sodium oxybate; Modafinil |
| 2021-046 | 83 | m | 442 | 6.9 | 1355 | 4 | A | 0 | Severe heart failure, died within 24 hrs after fall with hip fracture | Sodium oxybate, Paroxetine |
| **Narcolepsy type 1 with chronic opiates** |  |  |  |  |  |  |  |  |  |  |
| 2010-064 | 85 | f | 220 | 6.8 | 1206 | 1 | O | 0 | Chronic pain syndrome with palliative sedation | Morphine (9 years); Modafinil; Midazolam (last 24 hours) |
| **Control** |  |  |  |  |  |  |  |  |  |  |
| 2012-052 Hyp | 64 | f | 340 | 6.4 | 1221 | 0 | O | 0 | Legal euthanasia | Oxycodone, morphine, fentanyl, prednisolone, promethazine, codeine, oxazepam |
| 2000-022 Hyp LC | 83 | f | 470 | 6.5 | 1072 | 2 | O | 0 | Heart failure, Cachexia | - |
| 2012-005 Hyp | 84 | f | 336 | 6.7 | 1027 | 2 | A | 0 | Heart failure | Gabapentine, oxycodone, haloperidol |
| 2009-001 Hyp | 88 | m | 283 | 6.2 | 1418 | 2 | A | 0 | Gastro-intestinal bleeding | - |
| 1998-104 Hyp | 74 | f | 445 | 7.0 | 1167 | 2 | O | 0 | Cachexia in endstage pancreas carcinoma | Promethazine |
| 2018-005 Hyp | 76 | f | 330 | NA | 1030 | 1 | N | 1 | Respiratory insufficiency end-stage COPD | - |
| 2013-016 Hyp | 83 | m | 315 | 6.6 | 1372 | 1 | O | 1 | Myocardial infarction and palliative sedation | Oxazepam |
| 2012-094 Hyp SN | 82 | f | 315 | 6.3 | 1221 | 2 | B | NA | Thoracic aortic dissection | Levothyroxine, temazepam |
| 1998-036 SN | 69 | f | 375 | 6.6 | 1229 | 1 | NA | 0 | Cardiogenic shock | Levothyroxine, lorazepam |
| 2017-131 SN | 71 | f | 375 | NA | 1175 | 2 | NA | NA | Heart Failure | Duloxetine, fentanyl, codeine, pregabalin, quetiapine, temazepam, zopiclone |
| 2018-105 SN | 86 | m | 405 | 6.39 | 1340 | 3 | O | 0 | Intra-abdominal leakage of fluid causing abdominal infection | - |
| 2011-021 SN LC | 85 | f | 425 | NA | 1007 | 1 | B | 0 | Terminal renal insufficiency | Clonazepam, diazepam, amitriptyline, morphine, zopiclone, tramadol, pramipexol, fentanyl, midazolam |
| 2016-080 SN | 83 | m | 305 | 7.12 | 1108 | 2 | C | 2 | Legal Euthanasia | - |
| 2001-094 LC | 82 | f | 315 | 6.3 | 1221 | 2 | B | NA | Thoracic aortic dissection | Finasteride |

**Table 1 Clinical-pathological information**
|  |  |  |  |  |  |  |  |  |  |  |
| --- | --- | --- | --- | --- | --- | --- | --- | --- | --- | --- |
| 2001-139<br>LC | 73 | f | 815 | 6.30 | 1231 | 2 | C | 0 | Respiratory<br>insufficiency | Oxazepam,<br>paroxetine,<br>prednisolone,<br>haloperidol,<br>ketanserine |
| 2011-072<br>LC | 76 | f | 435 | 6.87 | 1072 | 2 | O | 0 | Hepatic failure and<br>colon carcinoma | Prednisolone,<br>codeine, fentanyl,<br>temazepam |
| 2017-093<br>LC | 82 | m | 345 | 7.28 | 1195 | 2 | A | 0 | Legal Euthanasia | Codeine,<br>prednisolone |

**Abbreviations:** Hyp, control for hypothalamus; SN, control for substantia nigra; LC, control for locus coeruleus ; ~, around that time; A, Amyloid; BW, Brain weight gram; B-AD, Braak Alzheimer's disease tangle stages (PMID,8307060); B-PD, Braak Parkinson's disease Lewy body's stages (PMID,12498954); f, female; m, male; NA, not available; NBB, Netherlands Brain Bank identification number; PMD, post mortem delay in minutes.

### Immunohistochemistry on human tissue

All formalin-fixed paraffin-embedded tissue blocks were coronally serially sectioned at 6 µm thick sections. The anatomical structures and borders including PVN, ME, Hcrt region, SN and LC were delineated by thionin and further confirmed by immunohistochemical staining (Shan *et al*., 2023).

In brief, after deparaffinization in xylene and rehydration through a graded ethanol series, postmortem human brain tissue sections were rinsed in distilled water. Endogenous peroxidase activity and non-specific binding was blocked by incubating the sections in a solution containing 3% H_2_O_2_ + 0.2% Triton X-100, and Tris buffered saline (TBS) 1x pH 7.6 for 30 minutes. This was followed by three washes in 1× TBS (pH 7.6), each for 5 minutes. The sections were then incubated in a blocking solution consisting of 10% normal horse serum in supermix (0.25g of gelatin, 0.5ml of Triton X-100 in 100 ml TBS) for 60 minutes at room temperature. Details of the antigen retrieval methods, as well as the specific primary and secondary antibodies used in this study, are provide in Supplementary Table 2, which includes antibody specificity and supporting references (PubMed ID). Tonsillitis tissue was included and served as positive control for all the CD4, CD8, CD79a staining or double staining.

### Quantification strategies

All quantifications were performed blinded to the nature of patients or controls by one person (M.F.). Microscopy protocols, software and parameters utilized in this study have been previously described (Shan *et al*., 2022, 2023). We previously delineated every areas on the basis of thionin and immunohistochemistry staining (Shan *et al*., 2022, 2023), then used immunohistochemical staining to count CD4^+^ or CD8^+^ cells. The perivascular space was identified based on the characteristic structure of blood vessels under light microscopy (Galiano-Landeira *et al*., 2020). T-cells within this compartment were counted as attached or closely associated with the vascular wall. In contrast, parenchymal T-cells were identified as being clearly separated from blood vessels.

Due to the scarcity of tissues, first we quantified T-cells in the immunohistochemically-identified peak sections—defined as the sections containing the highest density of CRH neurons, CRH fibers, Hcrt neurons, and TH-positive neurons in the PVN, ME, Hcrt, SN, and LC regions, respectively (Shan *et al*., 2022, 2023). When there was a positive finding, we further stained and systematically counted CD4^+^ and CD8^+^ T-cells throughout the PVN, ME, Hcrt regions at every 600 µm interval according to the Cavalieri’s principle described before (Shan *et al*., 2012). Moreover, to further confirm our findings, we repeated the CD4^+^ T-cells staining and counting throughout PVN, ME, Hcrt regions at every 600 µm interval.

The microglial markers IBA-1, HLA-DR, astrocyte marker GFAP, and blood-brain barrier leakage marker fibrinogen were positive for cell bodies and fibers. The integrated optical density (IOD) of IBA-1, HLA-DR and GFAP was quantified in peak sections containing the highest number of Hcrt neurons. The IOD of fibrinogen was measured in peak sections containing the highest number of Hcrt neurons, as well as in peak sections containing the highest number of immunocytochemically stained CRH neurons in PVN (Shan *et al*., 2022, 2023).

A series of fluorescence images were acquired using a 20x objective on the Axio Scan.Z1 slide scanner (Carl Zeiss GmbH, Jena, Germany), capturing signals from Alexa Fluor 488, Alexa Fluor 647, and DAPI channels. CD4^+^ T-cell profiles in the Hcrt region were visualized based on green fluorescence (Alexa 488), red fluorescence for CXCR6 (Alexa 647), and blue nuclear staining (DAPI), with image analysis performed using QuPath software (version 0.2.3). In a separate staining combination, CD4^+^ T-cells in the Hcrt region were labeled with red fluorescence (Alexa 647), green fluorescence for CD49a (Alexa 488), and DAPI. A third panel involved the detection of CD4^+^ or CD3^+^ T-cells in the Hcrt region, labeled with Alexa 647 and Alexa 488 respectively, along with DAPI for nuclear counterstaining. Single DAB-nikle staining for CD4^+^ T-cells resulted in minimal background (Figure 2). However, fluorescence-based detection in the human hypothalamus showed high background noise levels and strong lipofuscin autofluorescence. Since auto fluorescent and non-specific signals were negative for nuclear DAPI staining, only cells triple-labeled with CD4^+^ markers and DAPI were considered specific. These triple-positive cells were descriptively presented at the Hcrt level, which were close to the highest number of CD4^+^ T-cells observed in the brightfield-based systematic counts.

### Statistical analyses

NT1 and controls were compared using the exact Wilcoxon**-**Mann-Whitney U test (P). The false discovery rate was corrected by Benjamini-Hochberg corrections (q) for multiple testing of measures in the same cluster. The NT1 case with chronic opiate use was descriptively listed. Correlations were tested by Spearman’s correlation coefficient. All P values are two-sided. All q values < 0.05 (*), < 0.01 (**), < 0.001 (***) are considered as significant. Fold changes were calculated using the median values. Group dta are expressed as means ± 95% confidence intervals. Statistical analyses were carried out using SPSS Statistics version 25.0 (SPSS Inc, Chicago, IL). The figures were made using GraphPad Prism version 8.2 (GraphPad Software, San Diego, CA, USA).

## Results

### The total number of CD4^+^, not CD8^+^ T-cells increased in NT1 Hcrt region

Resident T-cells in the brain are thought to occur after initial antigen challenge and to remain lifelong (Mueller and Mackay, 2016; Smolders *et al*., 2018; Hsiao *et al*., 2023). We therefore counted the total number of CD4^+^ and CD8^+^ T-cells in the hypothalamic and brainstem tissues of NT1 brains and sex and age-matched. A significant 11-fold increase in the total number of CD4^+^ T-cells was found in NT1 patients compared to controls in the Hcrt area (P = 0.027*, q = 0.041*, Figure 1B), but not in PVN or ME (P ≥ 0.142, q ≥ 0.248 Figure 1A, C). The total number of CD8^+^ T-cells in control PVN, Hcrt region and ME were not different from those found in NT1 patients (P ≥ 0.149, q ≥ 0.447, Figure 1D-F). Furthermore, CD4^+^ T-cell staining, and quantification were repeated across the PVN, ME, and Hcrt regions, confirming the initial findings. In the NT1 individual and chronic use of opiates (NBB2010-064), total CD4^+^ (Figure 1A-C) or CD8^+^ (Figure 1D-F) cell number was in the same range as that of controls.

**Figure 1.**
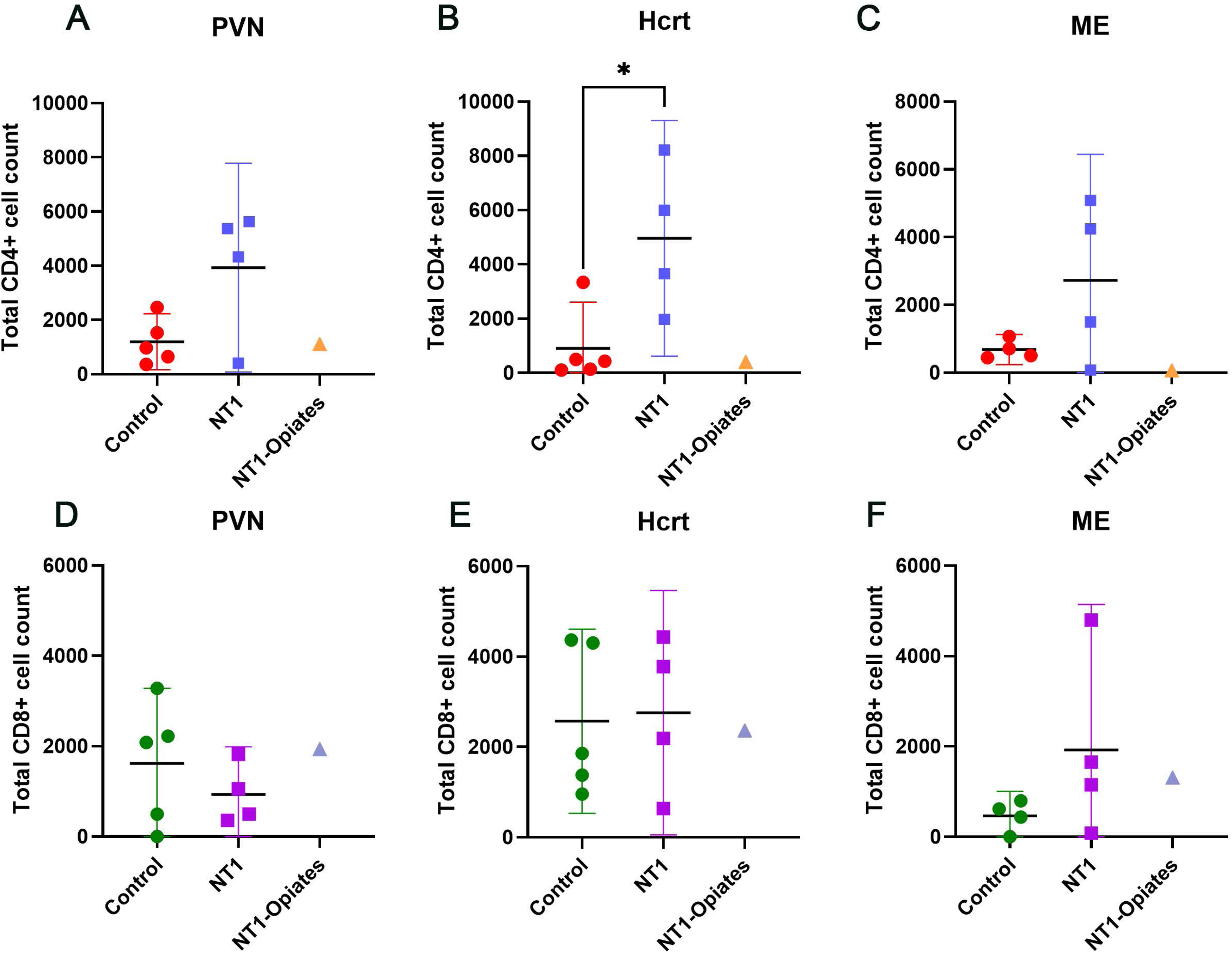
CD4^+^ T-cells but not CD8^+^ T-cells numbers are increased in the hypocretin (Hcrt) region of patients with narcolepsy type 1 (NT1). In the hypocretin region (Hcrt), the total number of CD4^+^ T-cells (B) was significantly increased in patients with NT1 compared to matched controls (P = 0.027*, q = 0.041*), but not in PVN (A) or median eminence (ME)(C). (D-F) The total number of CD8^+^ T-cells was unchanged in patients with NT1 compared to controls in PVN (D), Hcrt area (E) or ME (F). Bar plots showed mean and the lower Bound-Upper Bound of the 95% confidence intervals. The individual with NT1 and chronic opiate use (NT1 + opiates) is shown separately. The CD4^+^/CD8^+^ T-cell numbers of this NT1 + opiates case is relatively low, falling well within the range of controls.

### NT1 patients show an increased number of CD4^+^ T-cells but not CD8^+^ T-cells in the parenchymal and perivascular regions

Since the perivascular space has been identified as a niche for T-cells in the human brain (Smolders *et al*., 2013, 2018; Hsiao *et al*., 2023, 2025), we estimated a spatial association of T-cells with the extraluminal side of discernable vessels (‘perivascular space’) versus T-cells without this contact (‘parenchyma’) in the PVN, Hcrt area and ME. In NT1, within the Hcrt region, the total number of CD4^+^ T-cells was significantly increased compared to matched controls, both in the parenchyma (19-folds, P = 0.027*, q = 0.049*, Fig. 2G-I) and in perivascular space (7.8-folds, P = 0.049*, q = 0.049*, Fig. 2J-L). The total number of parenchymal or perivascular CD4^+^ T-cells in the PVN and ME was not different between NT1 and controls (P ≥ 0.142, q ≥ 0.221, Figure 2A-C and 2D-F; P ≥ 0.248, q ≥ 0.496, Figure 2M-O and 2P-R). In the NT1 individual with chronic use of opiates (NBB2010-064), total CD4^+^ T-cells number was similar in range to that of controls in both parenchyma (Fig. 2C, I, O) and perivascular (Fig. 2F, L, R) compartments. Furthermore, we repeated CD4^+^ T-cell staining and quantification across the PVN, Hcrt region, and ME, and confirmed that significant changes were specifically localized to the Hcrt region.

**Figure 2:**
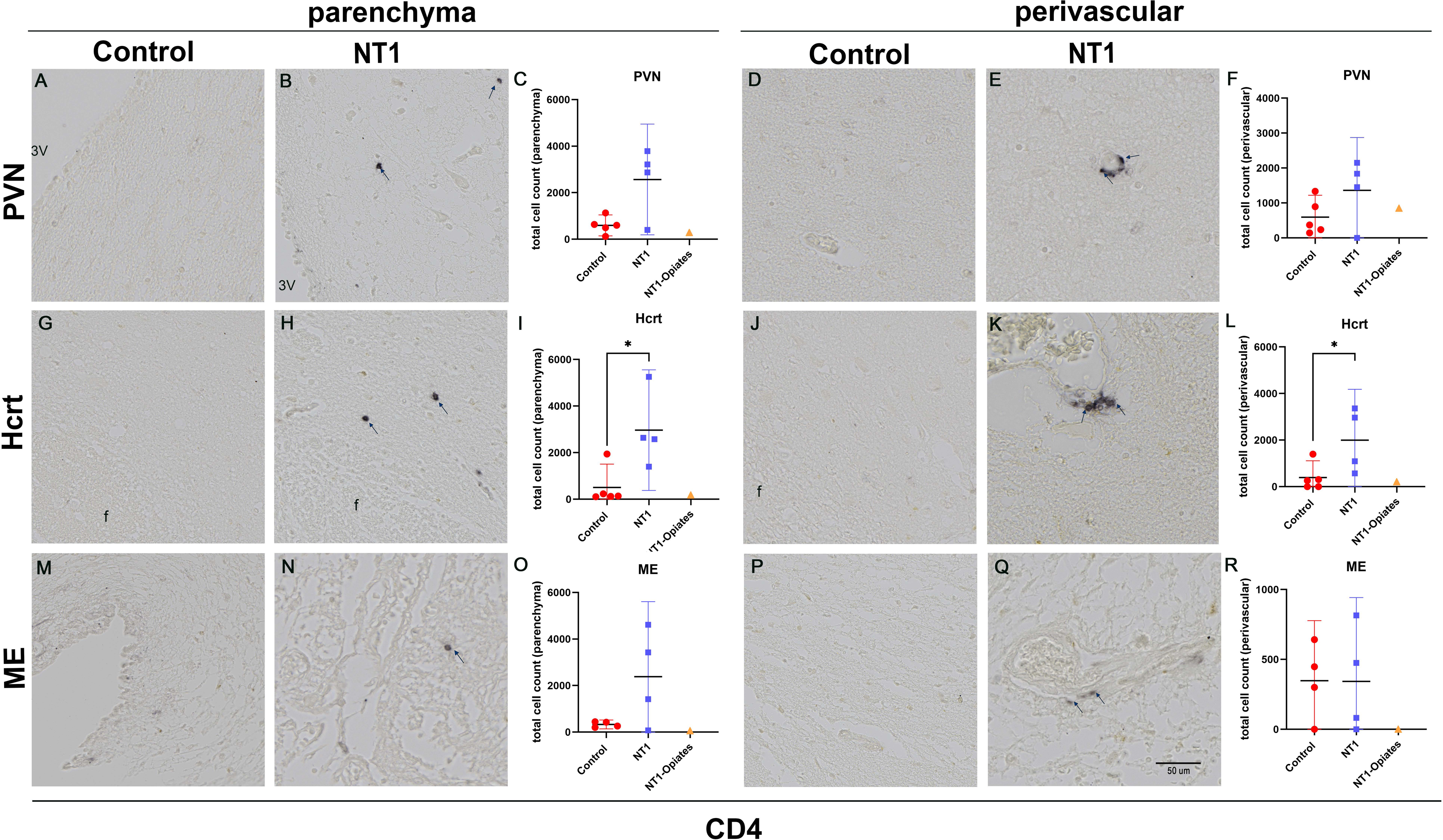
Increased number of CD4+ T cells in the parenchymal and perivascular spaces of the hypocretin (Hcrt) region of patients with narcolepsy type 1 (NT1). Representative images of the parenchymal CD4^+^ T-cells in the paraventricular nucleus (PVN) of a control (A) and a NT1 patient (B). The total number of parenchymal CD4^+^ T-cells (C) was unchanged in NT1 compared to controls. Representative images of the perivascular CD4^+^ T-cells in the paraventricular nucleus (PVN) of a control (D) and a NT1 patient (E). The total number of perivascular CD4^+^ T-cells (F) was unchanged in NT1 compared to controls. Representative images of the Hcrt region in a control (G) and a NT1 patient (H). The total number of parenchymal CD4^+^ T-cells (I) was significantly increased in patients with NT1 compared to controls (P = 0.027*, q = 0.049*). Representative images of the Hcrt region in a control (J) and a NT1 patient (K). The total number of perivascular CD4^+^ T-cells (L) was significantly increased in patients with NT1 compared to controls (P = 0.049*, q = 0.049*). Representative images of the parenchymal CD4^+^ T-cells in the median eminence (ME) in a control (M) and a NT1 (N). The total number of parenchymal CD4^+^ T-cells (O) was not different in NT1 compared to controls. Representative images of the perivascular CD4^+^ T-cells in the median eminence (ME) in a control (P) and a NT1 (Q). The total number of perivascular CD4^+^ T-cells (R) showed no difference in NT1 compared to controls. CD4^+^ T-cells were indicated by arrows. Bar plots in C, F, I and L, O, R showed mean and the lower Bound-Upper Bound of the 95% confidence intervals. The individual with NT1 and chronic opiate use (NT1 + opiates) is shown separately. The CD4^+^ T-cell numbers of this NT1 + opiates case is relatively low, falling well within the range of controls. f indicates the fornix and 3V is the region of third ventricle. Scale bar indicated 50 µm.

In contrast, the number of total parenchymal and perivascular CD8^+^ T cells in PVN, Hcrt and ME were not different between NT1 and controls ( P ≥ 0.081, q ≥ 0.447 ; parenchymal: Figure 3A-C, 3G-I and 3M-O, and perivascular: Figure 3D-F, 3J-L and 3P-R). The total CD8^+^ T-cells of NT1 individual with chronic use of opiates (NBB2010-064) fell well within the control group range, both in the parenchyma (Fig. 3C, I, O) perivascular compartments (Fig. 3F, L, R).

**Figure 3.**
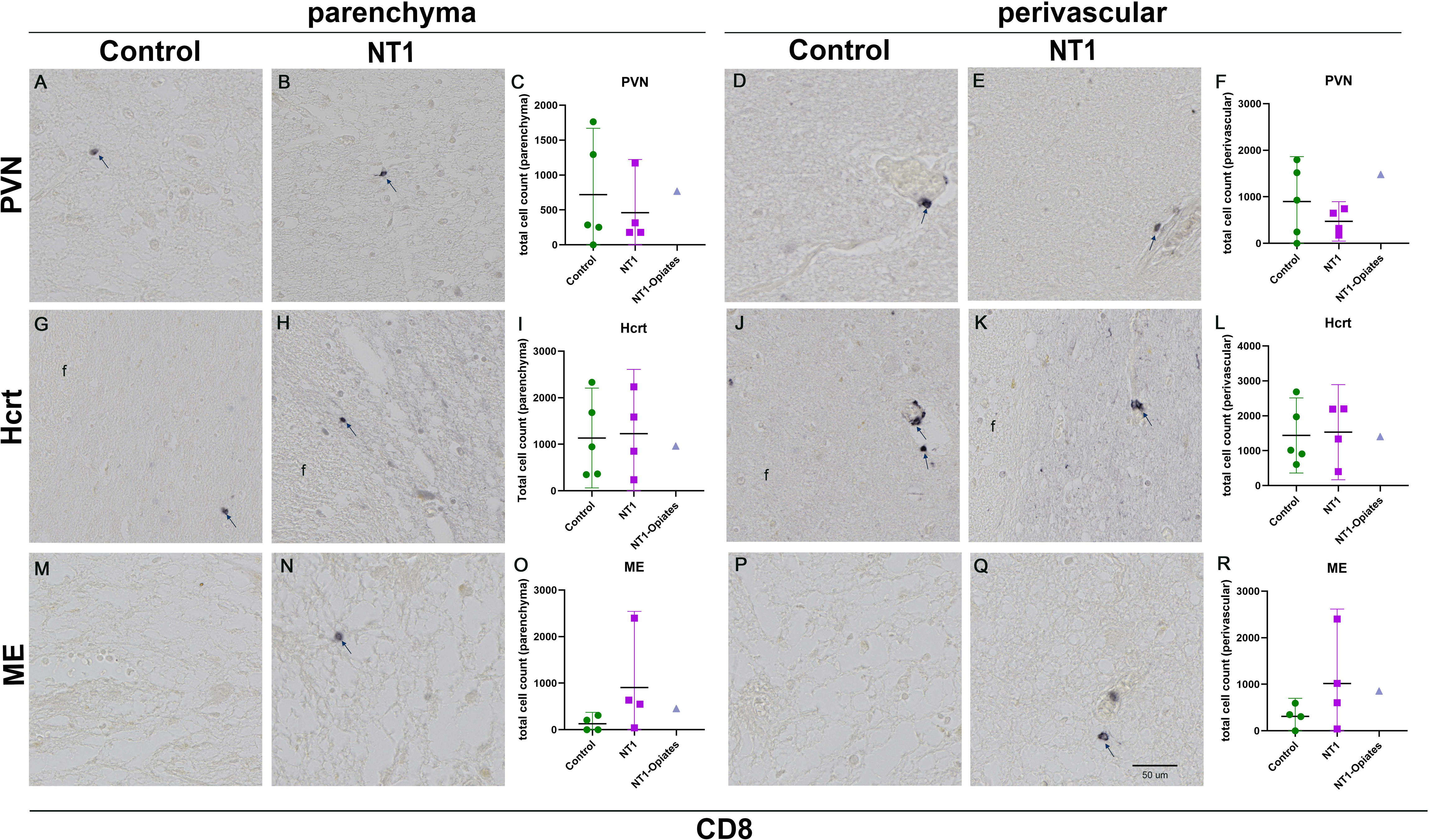
No difference in the number of parenchymal and perivascular CD8^+^ T-cells in the hypothalamic regions of patients with narcolepsy type 1 (NT1). Representative images of the parenchymal CD8^+^ T-cells in the paraventricular nucleus (PVN) of a control (A) and a NT1 patient (B). The total number of parenchymal CD8^+^ T-cells (C) was unchanged in NT1 compared to controls. Representative images of the perivascular CD8^+^ T-cells in the paraventricular nucleus (PVN) of a control (D) and a NT1 patient (E). The total number of perivascular CD8^+^ T-cells (F) was unchanged in NT1 compared to controls. Representative images of the Hcrt region in a control (G) and a NT1 patient (H). The total number of parenchymal CD8^+^ T-cells was unchanged in patients with NT1 compared to controls. Representative images of the Hcrt region in a control (J) and a NT1 patient (K). The total number of perivascular CD8^+^ T-cells (L) was unchanged in patients with NT1 compared to controls. Representative images of the parenchymal CD8^+^ T-cells in the ME in a control (M) and a NT1 (N). The total number of parenchymal CD8^+^ T-cells (O) showed no difference between NT1 and controls. Representative images of the perivascular CD8^+^ T-cells in the ME in a control (P) and a NT1 (Q). The total number of perivascular CD8^+^ T-cells (R) was not different in NT1 compared to controls. CD8^+^ T-cells were indicated by arrows. Bar plots in C, F, I and L, O, R showed mean and the lower Bound-Upper Bound of the 95% confidence intervals. The individuals with NT1 and chronic opiate use (NT1 + opiates) is shown separately. The CD8^+^ T-cell numbers of this NT1 + opiates case is relatively low, falling well within the range of controls. f indicated the fornix and 3V was the region of third ventricle. Scale bar indicated 50 µm.

**Figure 4.**
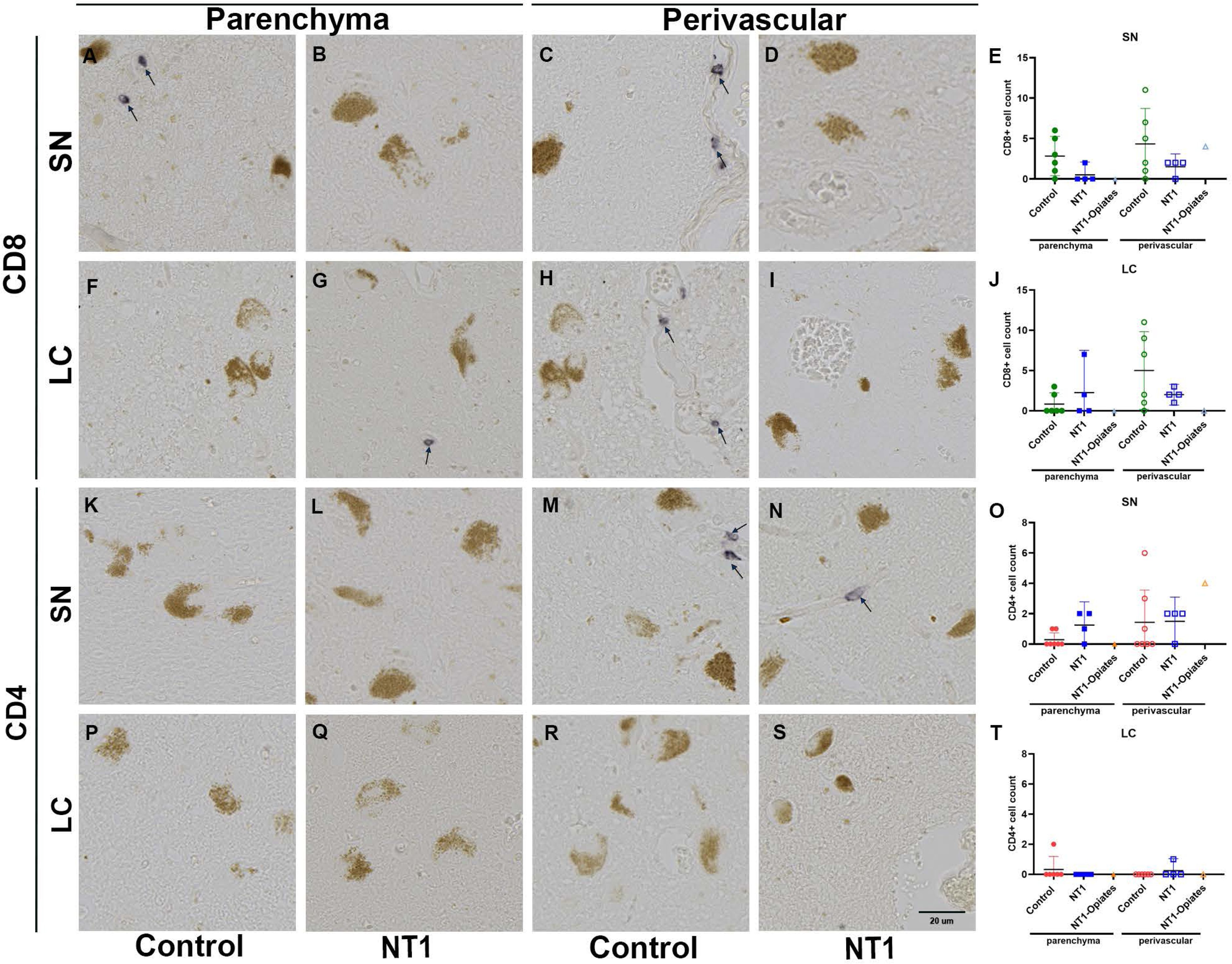
CD4^+^ and CD8^+^ T-cell numbers are unchanged in the Substantia Nigra (SN) and Locus Coeruleus (LC) of NT1 patients compared with controls. Representative images of parenchymal CD8□ T-cells in the Substantia Nigra (SN) of a control subject (A) and a NT1 patient (B), and perivascular CD8^+^ T-cells in the SN of a control (C) and a NT1 patient (D). Quantification of CD8^+^ T-cells in the SN (E) revealed no significant differences between NT1 patients and controls in either the parenchymal or perivascular compartments. Representative images of parenchymal CD8^+^ T-cells in the Locus Coeruleus (LC) of a control (F) and a NT1 patient (G), and perivascular CD8^+^ T-cells in the LC of a control (H) and a NT1 patient (I). Quantitative analysis in the LC (J) showed no significant difference in CD8^+^ T-cell numbers between groups in both compartments. Representative images of parenchymal CD4^+^ T-cells in the SN of a control subject (K) and a NT1 patient (L), and perivascular CD4^+^ T-cells in the SN of a control (M) and a NT1 patient (N). Quantification of CD4^+^ T-cells in the SN (O) revealed no significant differences between NT1 patients and controls in either the parenchymal or perivascular compartments. Representative images of parenchymal CD4^+^ T-cells in the Locus Coeruleus (LC) of a control (P) and a NT1 patient (Q), and perivascular CD4^+^ T-cells in the LC of a control (R) and a NT1 patient (S). Quantitative analysis in the LC (T) also showed no significant difference in CD4^+^ T cell numbers between groups across both compartments. CD8^+^/CD4^+^ T-cells were indicated by arrows. Bar plots in E, J, O and T showed mean and the lower Bound-Upper Bound of the 95% confidence intervals. The individual with NT1 and chronic opiate use (NT1 + opiates) is shown separately. The CD4^+^/CD8^+^ T-cell numbers of this NT1 + opiates case is relatively low, falling well within the range of controls. Scale bar indicated 20 µm.

Thus, we describe a localized enrichment of CD4^+^ T-cells in NT1 patients, specifically in the perivascular and parenchyma compartments of the Hcrt region, not seen in the other hypothalamic regions, such as the PVN and ME.

### No difference between groups in CD4^+^ and CD8^+^ T cell numbers in the SN and the LC

To establish whether NT1 changes are selective to the hypothalamus CD8^+^ and CD4^+^ T-cell numbers from peak sections the SN and LC were counted. There was no significant difference between NT1 patients and controls in either the perivascular space or parenchyma of the SN and LC (P ≥ 0.185; q ≥ 0.408 Figure 5A–T). The number of CD8^+^ and CD4^+^ T-cells of the individual with NT1 and chronic use of opiates (NBB2010-064) was similar in range to the controls the parenchyma and perivascular compartments. We conclude that the enrichment of CD4^+^ T-cells observed in NT1 patients is region-specific and localized to the hypothalamus, as it is not present in the SN and the LC.

**Figure 5.**
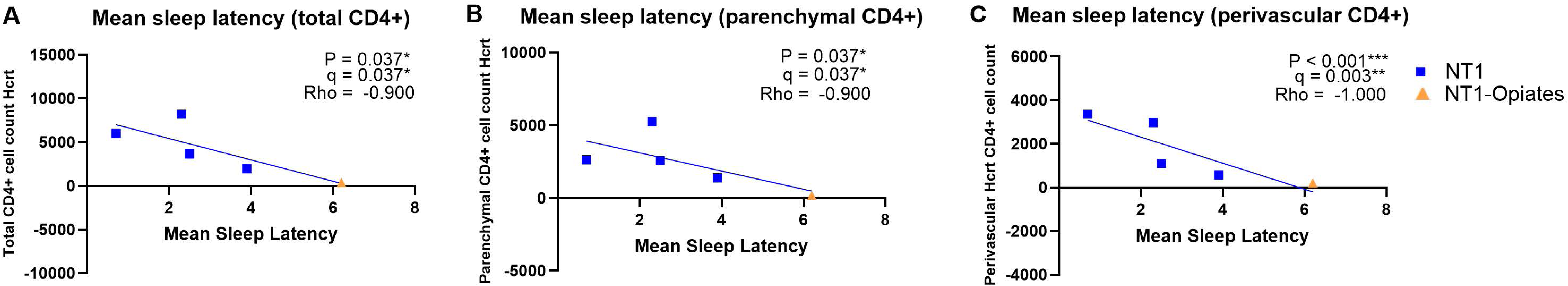
Significant negative correlation between CD4^+^ T-cell numbers and mean sleep latency in NT1patients. (A) The total number of CD4^+^ T-cells in the hypocretin (Hcrt) region shows a significant negative correlation with mean sleep latency in NT1 patients. (B, C). When stratified into parenchymal (B) and perivascular (C) compartments, CD4^+^ T-cell counts remained significantly negatively correlated with mean sleep latency. Trend lines in each scatter plot indicate the direction of the correlations.

### HLA DQB1*06:02 carriership in NT1 patients does not contribute to an overall lower ratio of CD8/CD4

To investigate whether HLA-DQB1*06:02 carriership in NT1 patients alone predisposes to a lower CD8/CD4 ratio compared to non-carrier controls, we compared CD8/CD4 ration in three different brain regions. Accordingly, the total CD8/CD4 ratio was significantly lower in hypothalamic of NT1 compared to non-carrier controls (P = 0.038*, q = 0.038* Supplementary Figure 1). No changes of CD8/CD4 ratio in SN or in LC were observed between NT1 and controls (P ≥ 0.136, q ≥ 0.136, Supplementary Figure 1). This may at least partly point to that the reduction of CD8/CD4 ratio in NT1 is a local hypothalamic-specific phenomenon, but not a donor-related trait due to HLA-DQB1*06:02.

### Phenotypic characterization of CD4^+^ T-cells in the Hcrt region suggests residential memory features

Brain CD4^+^ residential memory T-cells exhibited high expression of CD49a and CXCR6 mRNA and protein expression (Hsiao *et al*., 2023, 2025; Pignata *et al*., 2025). To establish whether the enriched CD4^+^ T-cell fraction shows residential memory potential, staining revealed that in two NT1 cases (NBB2018-091 and 2021-046), in the Hcrt region, CD4^+^ T-cells stained with DAPI were positive for CXCR6 (Figure 6A-D) or CD49a (Figure 6E-H), suggesting their identity as residential memory T-cells in the Hcrt region. In addition, to test whether the CD4^+^ staining observed was predominately due to T-cells but not macrophages or microglia, a triple staining using CD4^+^, CD3^+^ and DAPI was performed on the Hcrt region of two NT1 cases (NBB 2018-091 and NBB 2021-046) due to scarcity of tissue. All CD4^+^ T-cells (n=16), were positive to CD3^+^ and DAPI (Figure 6 I-L). The autofluorescence and nonspecific lipofuscin staining prevented further quantification. Thus, the identified enriched CD4^+^ cells were primarily T-cells, as shown by CD3^+^ and CD4^+^ reactivity, while co-staining with either CXCR6 or CD49a, identified characteristics of resident memory within the Hcrt region of NT1 patients.

**Figure 6.**
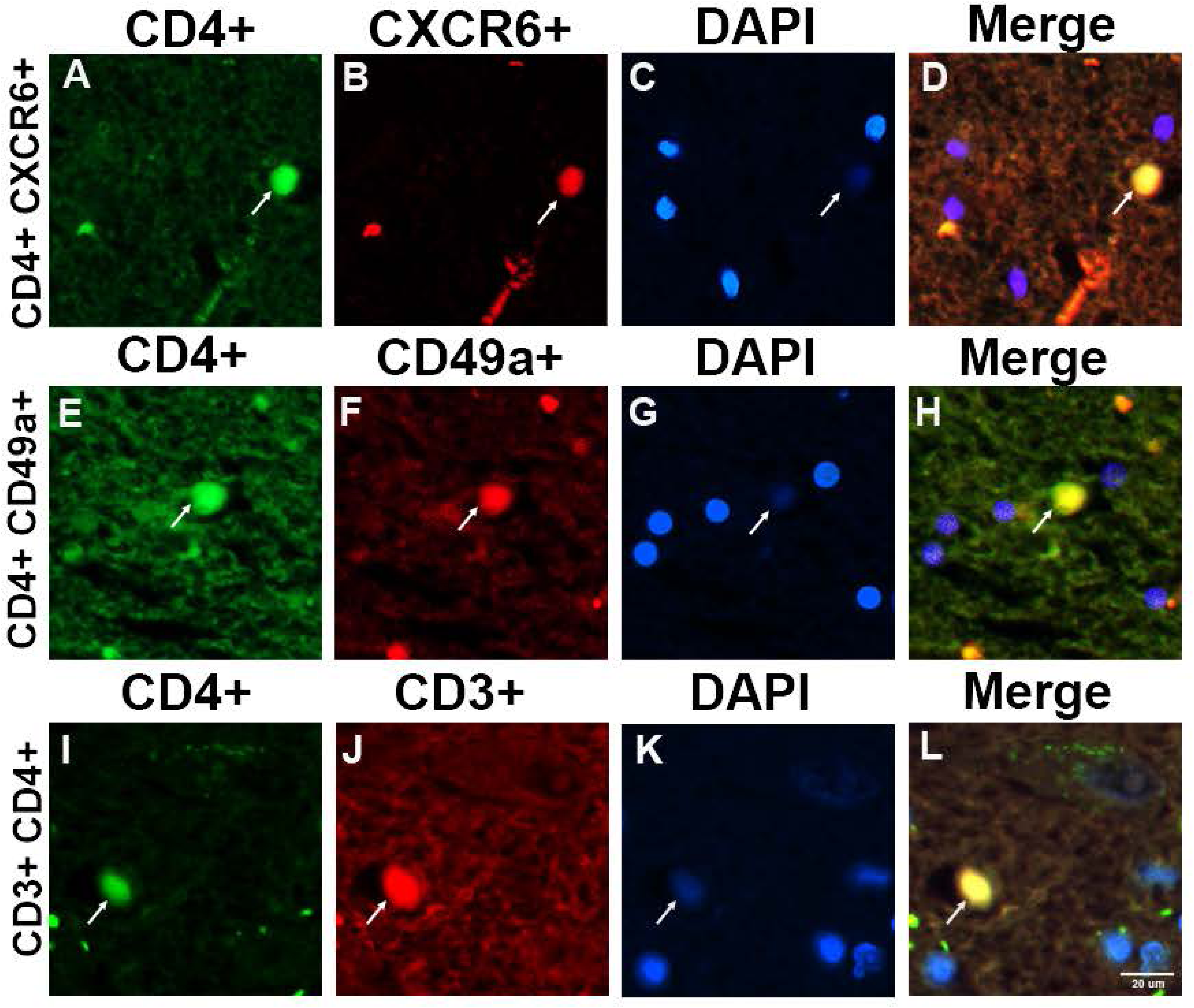
CXCR6 and CD49a were positive expressed with CD4^+^, denoting residential memory markers. (A, D) Representative immunofluorescence images of staining of CD4^+^ T-cells (A), CXCR6 positive (B), nuclei marked with DAPI (C) and all channels merged (D). The co expression of CD4 and CXCR6 indicates that a subset of CD4□ T-cells express tissue residential memory characteristics. (E, H) Representative immunofluorescence images of staining of CD4^+^ T-cells (E), CD49a positive (F), nuclei marked with DAPI (G) and all channels merged (H). The colocalization of CD4^+^ and CD49a indicates that a subset of CD4□ T-cells express tissue residential memory characteristics. (I,L) Representative immunofluorescence images of staining of CD4^+^ T-cell (I), CD3^+^ T-cell (J), nuclei marked with DAPI (K) and all channels merged (L). The colocalization of CD4^+^ and CD3^+^ indicates that CD4^+^ is expressed by T-cells. Scale bar indicated 20 µm.

### Hypothalamic IBA-1, HLA-DR, GFAP, and CD79a integrated optical density (IOD) does not differ between groups

The IOD of the microglial markers, IBA-1 in the Hcrt region, was not significantly different between control and NT1 cases ( P = 0.071, q = 0. 213, Table 2). In line with this observation the HLA-DR, -DP, -DQ staining, indicators of macrophage and microglia activation (Huitinga *et al*., 2004), was similar between NT1 and control cases in the Hcrt region ( P = 1.000, q = 1.000, Table 2). In addition, there is no difference between groups in IOD for the astrocyte marker GFAP in the Hcrt region ( P = 0.730, q = 1.000, Table 2).

**Table 2.** Summary of Immunocytochemistry quantification.

| Structure | Primary antibody | Q | NT1 | Control | NT1(opiates) | R(95%CI) |
| --- | --- | --- | --- | --- | --- | --- |
| Hcrt | Iba1 | IOD | 0.009±0.003 | 0.001±0.001 | 0.006 | 0.111 [0.006 0.217] |
| Hcrt | HLA-DR | IOD | 0.027±0.014 | 0.038±0.016 | 0.005 | 1.395 [0.02 1.942] |
| Hcrt | GFAP | IOD | 0.033±0.008 | 0.030±0.007 | 0.047 | 0.909 [0.046 1.772] |
| PVN/ Hcrt | Fibrinogen | IOD | PVN (0.001±0.001)<br>Hcrt (0.032±0.011) | PVN (0.011±0.0031)<br>Hcrt (0.055±0.015) | PVN (0)<br>Hcrt (0.085±0.017) | PVN (11.00 [0.550 21.450])<br>Hcrt (1.620 [0.081 3.159]) |
| PVN | CD4+ | CC | total(3930.375±1209.476)<br>par (2569.375±746.907)<br>periv(1361.000±475.653) | total (1194.600±372.603)<br>par (598.100±161.598)<br>periv (596.400±225.328) | total (1132.000)<br>par (285.000)<br>periv (847.000) | total (0.304 [0.015 0.593])<br>par (0.233 [-0.590 0.291])<br>periv(0.438 [0.022 0.854]) |
| Hcrt | CD4+ | CC | total(4965.000±1363.879)<br>par (2967.000±814.443)<br>periv (1998.000±686.024) | total (901.000±614.850)<br>par (508.400±359.593)<br>periv (392.600±225.328) | total (384.500)<br>par(177.500)<br>periv (207.000) | total (0.222 [0.01 0.447])<br>par (0.171 [-1.599 0.177])<br>periv (0.196 [0.01 0.382]) |
| ME | CD4+ | CC | Total(2726.650±1168.656)<br>par (2383.25±1012.498)<br>periv (343.400±188.410) | total(681.750±140.722)<br>par (334.25±59.770)<br>periv (347.600±2135.352) | total (57)<br>par (57.000)<br>periv (0) | total (0.259 [0.01 0.505])<br>par (0.140 [0.007 0.273])<br>periv(1.145 [0.065 2.225]) |
| PVN | CD8+ | CC | total (933.751±331.835)<br>par (462.5±239.075)<br>periv (472.500±123.345) | total (1616.120±600.572)<br>par (718.6±342.190)<br>periv (897.600±348.052) | total (1636.0)<br>par (160.0)<br>periv (1476) | total (1.731 [0.087 3.375])<br>par (1.554 [0.080 3.02])<br>periv(1.900 [0.095 3.705]) |
| Hcrt | CD8+ | CC | total (2758±849.541)<br>par (1227.5±868.27)<br>periv(1530.500±427.896) | total (2571.000± 733.794)<br>par (1134.2±862.98)<br>periv (1437.200± 386.795) | total (2356)<br>par (958.000)<br>periv (1398.000) | total (0.933 [0.047 1.819])<br>par(0.924 [0.047 1.801])<br>periv(0.587 [0.030 1.144]) |
| ME | CD8+ | CC | total (1921.500±1021.54)<br>par (905.5±513.770)<br>periv (1016.000±502.844) | total (440.500±173.539)<br>par (128.500±77.167)<br>periv (312.000±121.693) | total (1300.00)<br>par (450.000)<br>periv (850.000) | total (0.241 [0.012 0.470])<br>par(0.142 [0.007 0.277])<br>periv(0.307 [0.016 0.598]) |
| SN | CD4+ | CC | total (3.000±0.707)<br>par (1.250±0.478)<br>periv (2.000±0.577) | total (2.857±1.639)<br>par (0.571±0.202)<br>periv (2.286±1.523) | total (2.000)<br>par(2.000)<br>periv (0) | total (0.952 [0.048 0.1856])<br>par (0.457 [0.023 0.891])<br>periv(1.143 [0.057 2.229]) |
| LC | CD4+ | CC | total (1.000±0.408)<br>par (0.500±0.289)<br>periv 0.500±0.224) | total (0.833±0.543)<br>par (0.333±0.211)<br>periv (0.500±0.315) | total (2.000)<br>par (1.000)<br>periv (1.000) | total (0.833 [0.042 1.624])<br>par (0.667 [0.034 1.300])<br>periv (1.000 [0.050 1.950]) |
| SN | CD8+ | CC | total (5.000±1.633)<br>par (0.750±0.478)<br>periv (4.250±1.702) | total (9.000±3.579)<br>par (2.429±0.870)<br>periv (6.571±2.793) | total (0)<br>par (0)<br>periv (0) | total (1.800 [0.090 3.510])<br>par (3,239 [0.162 6,316])<br>periv (0.180 [0.009 0.351]) |

**Table 2 Summary of Immunocytochemistry quantification**
|  |  |  |  |  |  |  |
| --- | --- | --- | --- | --- | --- | --- |
| LC | CD8+ | CC | total (3.000±0.707)<br>par (1.250± 0.750)<br>periv (1.750±0.250) | total (4.50±1.648)<br>par (1.000± 0.471)<br>periv (3.500± 1.875) | total (2.000)<br>par (2.000)<br>periv (0) | total (1.500 [0.0752.925])<br>par (0.800 [0.040 1.560])<br>periv (1.400 [0.070 2.730]) |

**Abbreviations:**
95% CI: 95% confidence interval (lower bound, upper bound); CC: Cell Count; CD4<sup>+</sup>: Cluster of Differentiation 4 positive T-cells; CD8<sup>+</sup>: Cluster of Differentiation 8 positive T-cells; Fibrinogen: Plasma protein involved in clotting and inflammation (used as a Blood Brain Barrier marker here); GFAP: Glial Fibrillary Acidic Protein; Hcrt: Hypocretin-1 / Orexin A; also refers to medial lateral hypothalamus/hypocretin region; HLA-DR: Human Leukocyte Antigen-DR (immune-related microglia marker); IBA-1 (Iba1): Ionized calcium binding adaptor molecule 1 (microglia marker); IOD: Integrated Optical Density; LC: Locus Coeruleus; ME: Median Eminence; NT1: Narcolepsy Type 1 (narcolepsy with cataplexy); NT1 with opiates: Narcolepsy Type 1 patient who received opiates (NBB2010-064); NBB: Netherlands Brain Bank; par: Parenchyma; periv: Perivascular; PVN: Paraventricular Nucleus; Q: Quantification; R: Mean of ratio (Controls/NT1 mean); S.E.M.: Standard Error of the Mean; SN: Substantia Nigra

Since a leaky blood-brain barrier could enable migration of circulating T-cells into the brain, we stained for fibrinogen to estimate blood-brain barrier integrity within the Hcrt and PVN regions. Immunostaining for fibrinogen did not reveal differences in these regions for NT1 patients as compared to controls ( P ≥ 0.289, q ≥ 0.434 Table 2).

Since chronic perivascular inflammatory infiltrates are often composed of B-cells in inflammatory diseases, immunostaining for the B-cell marker CD79a was performed on two NT1 cases (with highest and lowest total Hcrt neuron number, NBB 2010-064 and NBB 2021-046), due to scarcity of tissue. No CD79a positive cells were detected in the PVN or hypocretin neuron regions in either case, despite robust positive staining in positive control tonsil tissue. Based on these negative findings, further investigation of this marker was discontinued. We conclude that perturbations in CD4^+^ T-cells are the most prominent changes in immune-subsets in NT1 Hcrt neuron regions.

### CD4^+^ T-cell number negatively correlated with mean sleep latency in NT1 patients

To investigate a potential association between CD4^+^ or CD8^+^ T-cell numbers and antemortem sleepiness in NT1 patients, Spearman correlation analyses were performed. A significant negative correlation was found between the total number of CD4^+^ T-cells in the Hcrt region and the mean sleep latency in NT1 cases (rho = -0.900, P = 0.037*, q = 0.037*, Fig. 5A). Further stratification revealed that both parenchymal CD4^+^ T cell counts (rho = -0.900, P = 0.037*, q = 0.037*, Fig. 5B) and perivascular CD4^+^ T-cell counts (rho = -1.000, P < 0.001***, q = 0.003**, Fig. 5C) were significantly negatively correlated with mean sleep latency. No such association was observed for CD4^+^ T-cell in PVN or ME regions or CD8^+^ T cells in three hypothalamic regions. This pointed out that shorter sleep latency is associated with higher number of CD4^+^ T-cells within Hcrt region.

## Discussion

We found an 11-fold increase in total CD4^+^ T-cell number in the Hcrt region of NT1 patients, as well as an 19-fold increased in the parenchymal region and 7.8-fold increase in the perivascular compartments, CD8^+^ T-cell numbers remained unchanged in all areas investigated. In addition, the increase in CD4^+^ T-cells was observed specifically in the Hcrt region of NT1 patients, but not in the PVN, ME, SN nor the LC. Moreover, we showed that since those CD4^+^ T-cells co-stained CXCR6 or CD49a, they displayed feature of resident memory T-cell in the Hcrt region of NT1 patients. In addition, the number of CD4^+^ T-cells in the Hcrt neuron region showed a negative correlation with mean sleep latency in NT1 cases. This points out that greater symptom severity—reflected by shorter sleep latency—is associated with higher number of CD4^+^ T-cells in this region. These cells are hypothesized to reflect the remnants of CD4^+^ T-cells which infiltrate the hypothalamus in NT1 patients and play a direct role in the autoimmune process targeting Hcrt neurons.

Our immunohistochemical finding showing a mean CD8/CD4 ratio in the range of 1.94-2.89 in control hypothalami, was in good agreement with the ratio estimated with staining and flow-cytometry in a previous study (Smolders *et al*., 2018). NT1 has long been suspected to have an immune-mediated pathogenesis given its tight association with HLA class II allele DQB1*06:02, mutations in T-cell receptor genes and other immune-relevant loci (JUJI *et al*., 1988; Mignot *et al*., 2001; Hallmayer *et al*., 2009; Han *et al*., 2014; Ollila *et al*., 2023). The increased incidence of NT1 that has been observed after H1N1 influenza infections and certain vaccines (Han *et al*., 2011; Stowe *et al*., 2020; Ollila *et al*., 2023) further substantiates this hypothesis. Recently, Hcrt autoreactive CD4^+^ and CD8^+^ T-cells were detected in the blood and CSF of NT1 patients (Latorre *et al*., 2018; Luo *et al*., 2018; Cogswell *et al*., 2019; Jiang *et al*., 2019; Pedersen *et al*., 2019). However, this may be caused by a secondary interaction with Hcrt that is released into circulation following the loss of Hcrt neurons (Liblau, 2018). Our results from postmortem human brain samples show for the first time that T-cells, especially CD4^+^ T-cells are selectively enriched in the Hcrt region in NT1 patients, as postulated by dominant HLA class II association with this disease. In contrast, there are no differences in CD8^+^ T-cell numbers between NT1 patients and controls. A case report showed an anti-Ma-associated encephalitis patients with NT1 symptoms with an associated CD8^+^ inflammatory-mediated response against Hcrt neurons (Dauvilliers *et al*., 2013). Our finding may indicate a more specific and selective autoimmune process targeting Hcrt neurons in NT1 patients.

While in NT1 animal model or in other neurological disease like Parkinson’s disease, the CD8^+^ components has been widely described, our current findings imply that in NT1 there is a more pronounced CD4^+^ component. In an NT1 animal model, a predominantly CD8^+^ T-cell-mediated cytotoxic reaction was observed following influenza vaccination or expression of hemagglutinin, a surface protein of influenza viruses, leading to the death of Hcrt neurons (Bernard-Valnet *et al*., 2016, 2022). Moreover, other neurological diseases such as Parkinson’s disease (Sulzer *et al*., 2017; Galiano-Landeira *et al*., 2020) are associated with increasing of CD8^+^ T-cells than CD4^+^ T-cells in the brain. CD8^+^ T-cells recognize antigens bound to HLA class I protein which can present on neurons. However, it remains unclear in NT1 how CD4^+^ T-cells target Hcrt neurons, because neurons do not express HLA class II proteins (Liblau *et al*., 2013). An interaction with transiently infiltrating B cells or myeloid cells in the acute phase can be postulated. Although low numbers of perivascular CD4^+^ T-cells are always encountered (Smolders *et al*., 2018; Hsiao *et al*., 2023, 2025), B-cells are only incidentally found in the post-mortem human brain in the absence of acute or ongoing chronic neuroinflammation (Fransen *et al*., 2021; Bogers *et al*., 2023). Alternatively, it may be postulated that the remaining CD4^+^ T-cells we observed were remnants of antigen spreading due to Hcrt cell loss. In the NT1 Hcrt region, the interactions of CD4^+^, CD8^+^ T-cells and other immune cells or antibodies possibly mediating this process deserve further investigations using spatial in situ RNA and epigenetic sequencing (Seifinejad *et al*., 2023). Interestingly, we observed a negative correlation between CD4^+^ T-cells in the Hcrt region and the antemortem mean sleep latency in NT1 cases, which is direct evidence that the NT1 disease severity is directly associated with the abundance of CD4^+^ T-cells in the Hcrt region.

It was expected that the total number of CD4^+^ T-cells would relatively small, because we determined the fraction of CD4^+^ T-cells of long-living tissue residence memory cells that persist decades beyond pathogen clearance at sites of antigenic challenge (Smolders *et al*., 2018; Künzli and Masopust, 2023). In our study, all four patients with typical NT1 had a history of disease from 46 to 68 years. We showed that the data cannot be explained by blood brain barrier leakage, B-cells, microglia or astrocyte activation and due to the time involved it is not likely to detect cytotoxic factors such as granzymes. Most neuroinflammation related cells are short-lived after disease onset. This is in line with the observation that higher percentage of CD4^+^ and CD8^+^ T-cells release cytokines in response to Hcrt in NT1 children who, as a group, were closer to NT1 onset compared to their adult patients (Cogswell *et al*., 2019). We did not find microglia changes in agreement with observations of others in postmortem brains of NT1 cases (Honda *et al*., 2009) and in PET scans of living patients (Barateau *et al*., 2024). Microglia alterations have been observed in a NT1 animal model, during the acute phase, where microglia became activated and proliferated around the region of Hcrt neuron degeneration (Tabuchi *et al*., 2014). However, by four weeks after the loss of these neurons, the microglia returned to a resting state (Tabuchi *et al*., 2014). In addition, CRH neurons in the PVN were less affected by T-cells as the current study didn’t observe a significant changes of T-cells in the PVN or the ME, yet clearly suggestive of higher levels deserves future study with a larger sample size. Together with the low and non-significant number of T-cells we found in the SN and LC this indicate that the CD4^+^ T-cell infiltration in NT1 hypothalamic is a Hcrt region selective process.

The study has several limitations. The first is that this is a descriptive and retrospective study and the etiopathogenetic relevance of CD4^+^ T-cell should be confirmed with brain samples from an earlier stage of disease that allows to investigate in the course of Hcrt neuronal loss and its relationship with CD4^+^ T-cells. Due to the rapid development of NT1, usually when the patients reached the clinic, they already show full clinical symptoms and a completely loss of Hcrt (Liblau *et al*., 2024). However, the individual with NT1 and chronic use of opiates (NBB2010-064) showed a 33% of control total number of Hcrt neurons. The CD4^+^ T-cell numbers of this case is relatively low, falling well within the range of controls. This might indicate a reduced infiltration of CD4^+^ T-cells which may correlate with a preservation of Hcrt neuronal numbers in NT1. Another limitation of the current study is the relatively small sample size, which limits power to detect differences between NT1 patients and controls and between other hypothalamic regions such as PVN and ME. Given that NT1 is a very rare disorder, postmortem brain tissue from NT1 patients is scarce. Nevertheless, even with five NT1 and five control cases, we were able to confirm the marked reduction in Hcrt neuronal numbers and increased in histaminergic neuronal numbers previously reported in the literature (Peyron *et al*., 2000; Thannickal *et al*., 2000; John *et al*., 2013; Valko *et al*., 2013). In addition, differentiating perivascular and parenchymal T-cells using endothelial or laminin staining warrants further investigation. However, the main conclusion was not affected, irrespective of whether CD4^+^ T-cells were analyzed as a total population or separated into perivascular and parenchymal compartments.

In conclusion, to the best of our knowledge, this is the first study to show an increased in CD4^+^ T-cells numbers, but not CD8^+^ T-cells in the Hcrt region of NT1 postmortem brains. Furthermore, we showed that CD4^+^ T-cells in Hcrt of NT1 patients, exhibit resident memory T-cells features. This helps to bridge -at least partly-the gap in the autoimmune hypothesis of NT1 by linking genetic T-cell related risk factors (JUJI *et al*., 1988; Mignot *et al*., 2001; Hallmayer *et al*., 2009; Han *et al*., 2014; Ollila *et al*., 2023) with observed increased in T-cell reactivity against Hcrt in the blood and CSF of NT1 patients (Latorre *et al*., 2018; Luo *et al*., 2018; Cogswell *et al*., 2019; Jiang *et al*., 2019; Pedersen *et al*., 2019). Our data suggest a primary role for CD4^+^ T-cells infiltration in Hcrt neuronal destruction. Furthermore, the negative correlation between CD4^+^ T-cells in the Hcrt region and shorter sleep latency suggests a role of these T-cells in the disease severity. Currently, NT1 diagnosis and treatment are focusing on late disease stages, i.e. after Hcrt destruction (Mignot *et al*., 2002; Bassetti *et al*., 2019). These findings further support the autoimmune etiology of NT1 and highlight the potential for T cell–based early diagnosis and immunomodulatory interventions in its treatment (Liblau *et al*., 2024).

## Acknowledge

This work was supported by a grant of the Friends Foundation of the Netherlands Institute of Neuroscience (Stichting Vrienden van het Herseninstituut). The authors are grateful to Prof. Inge Huitinga, director of the Netherlands Brain Bank (NBB) of the Netherlands Institute for Neuroscience for providing valuable brain samples, clinical and neuropathological information. The authors wishes to express their gratitude to experienced neurologists/somnologists (Prof. G. J. Lammers and Dr. R. Fronczek), who diagnosed and treated those NT1 patients. The authors are greatly indebted to Ms. A. L. Ciobanu Mr. J. Engelenburg, D. Wever and C. Hsiao for valuable suggestions as well as positive control tonsil samples for CD4/CD8 staining.

## Author Contributions

LS, and DFS contributed to the conception and design of the study. JS contributed to the conception and design of T-cell studies. LS, DFS and MF, MH contributed to the acquisition and analysis of data. MF, LS, and DFS contributed to drafting the text and figures.

## Data Availability Statement

Raw data were generated in the Netherlands Institute For Neuroscience. Derived data are available from the corresponding author on request.

## Legends

**Supplementary Figure 1. The CD8/CD4 ratio was significantly lower in hypothalamus of NT-1 patients compared to controls** (P = 0.038*, q = 0.038*). No changes of CD8/CD4 ratio in the SN or the LC was observed between NT1 and controls (P ≥ 0.136, q ≥ 0.136). Bar plots showed mean and the lower Bound-Upper Bound of the 95% confidence intervals.

