## Supplementary Table 1 for "Increased numbers of CD4^+^ T-cells in the hypocretin/orexin region of Narcolepsy Type 1"

| **NBB** | **Hcrt-CSF** | **Cataplexy** | **Additional diagnostic tests** |
| --- | --- | --- | --- |
| **Narcolepsy type 1** | | |  |
| 2008-023 | UD | Yes | MSLT(1998): 5 out of 5 sleep, mean sleep latency 0.7m and 3 SOREMs, HLA DQB1*0602 + |
| 2018-018 | UD | Yes | MSLT(1998 with Sodium oxybate): 5 out of 5 sleep; mean sleep latency 3.9m, no SOREMs. AHI 5.4, HLA DQB1*0602 + |
| 2018-091 | UD | Yes | MSLT(2017): mean sleep latency 2.3m, 3 SOREMs; PSG (2014): No clinically relevant sleep apnea; PSG (2017): AHI 39 (after this CPAP was started), HLA DQB1*0602 + |
| 2021-046 | NA | Yes | MSLT(2017): mean sleep latency 2.5m without SOREMs, PSG (2018). Diagnosis made in 1982 in Germany.: elevated AHI 47 under treatment with Sodium oxybate and paroxetine; elevated AHI considered not to be clinically relevant. HLA DQB1*0602 + |
| **Narcolepsy type 1 with chronic opiates** | | | |
| 2010-064 | 200 | Yes^  (age:61) | 1986: uncontrollable urge to sleep, typical cataplexy and sleep paralysis; PSG (1997): No sleep apnea.  MSLT (1998 without opiates, with imipramine): 5 out of 5 sleep, mean sleep latency 6.2m and 1 SOREM. HLA DQB1*0602 + |
| **Control hypothalamus** | | | |
| 2012-052 |  |  | HLA DQB1*0602 - |
| 2000-022 |  |  | HLA DQB1*0602 - |
| 2012-005 |  |  | HLA DQB1*0602 - |
| 2009-001 |  |  | HLA DQB1*0602 N.A. |
| 1998-104 |  |  | HLA DQB1*0602 N.A. |
| 2018-005 |  |  | HLA DQB1*0602 N.A. |
| 2013-016 |  |  | HLA DQB1*0602 N.A. |

**Abbreviations:** ^: cataplexy disappeared after taking opiates for chronic pain; +: positive; -: negative; ~: around that time; AHI: Apnea-hypopnea index; CPAP: continuous positive airway pressure machine; Hcrt-CSF: hypocretin (orexin) CSF concentration (pg/ml); Hypocretin-1 levels were measured in the Leiden University Medical Center with radioimmunoassay (RIA) (Phoenix Pharmaceuticals, Belmont, USA).; MSLT: multi sleep latency test; NA: not available; NBB: Netherlands Brain Bank identification number; PLMI: periodic leg movement index; PSG: Polysomnography; SOREMs: sleep onset rapid-eye-movement episodes; UD: undetectable levels.
