## Supplementary Table 2 for "Increased numbers of CD4^+^ T-cells in the hypocretin/orexin region of Narcolepsy Type 1"

| **Structure** | **Target, Primary antibody** | **Source, host species, Cat#., RRID** | **Specificity and characterization**  **(PMID)** | **Antigen retrieval and dilution** |
| --- | --- | --- | --- | --- |
| T-cell | CD4+ T-cells | Abcam, Rabbit, polyclonal, Cat# ab133616, **RRID**:AB_2750883 | (31080441) | MW, sodium citrate buffer (pH 6.0); 1:500 in Sumi-Natural Horse Serum |
| T-cell | CD4+ T-cells | R&D systems, Goat, polyclonal, Cat#AF-379-NA, Lot: YS1023101, RRID:AB_354469 | (37604932) | MW, sodium citrate buffer (pH 6.0); 1:100 in Sumi-Natural Horse Serum |
| T-cell | CD8+ T-cells | Abcam, Rabbit, monoclonal,Cat#ab4055, RRID:AB 304247 | (38136261) | MW, sodium citrate buffer (pH 6.0); 1:500 in Sumi-Natural Horse Serum |
| BBB | Fibrinogen | Dako, Rabbit, Polyclonal, Cat#A0080, | (31753008) | MW, sodium citrate buffer (pH 6.0), 1:7500 in Sumi |
| B-cell | CD79a | Dako, Mouse, Monoclonal, Cat# M705001-2, Clone JCB117 | (9924428) | MW, Tris-EDTA (pH 9.0), 1:200 in Sumi-Natural Horse Serum |
| T resident memory cells | CXCR6 | Abcam, Rabbit, Polyclonal, Cat#ab8023, Lot 104494-3 RRID:AB_306205 | (33829065) | MW, sodium citrate buffer (pH 6.0); 1:1000 in Sumi-Natural Horse Serum |
| T resident memory cells | CD49a (ITGA1) | Atlas Antibodies, Mouse, monoclonal, Cat# AMAb91461, Clone: CL7217 RRID:AB_2732111 |  | MW, sodium citrate buffer (pH 6.0); 1:1000 in Sumi-Natural Horse Serum |
| **Secondary antibody** |  |  |  |  |
|  | Alexa Fluor 647 conjugated Donkey Anti-Rabbit IgG (H+L) | Jackson Immunoresearch Labratories, Polyclonal, 711-606-152, RRID: AB_2340586, |  | 1:200 |
|  | Alexa 488 conjugated Donkey Anti-Goat IgG (H+L) | Jackson Immunoresearch Labratories, Polyclonal, 705-006-147 RRID: AB_2340386 |  | 1:200 |
|  | Alexa Fluor 488 Conjugated Donkey Anti-Rat IgG (H+L) | Jackson Immunoresearch Labratories, Polyclonal, Cat# 712-546-150, RRID:AB_2340685 |  | 1:200 |
|  | Goat anti Rabbit (BA-1000-1.5) | Vector Labratories Inc, Biotinylated . Cat# BA-1300, **RRID**:AB_2336188 |  | 1:400 |
|  | Alexa Fluor Goat Anti-Mouse (Alexa 488) | Jackson Immunoresearch Labratories, Polyclonal, 205-005-108 RRID: AB_2339054 |  | 1:200 |
| **Avidin-biotin complex** | ABC Elite kit | Vector Laboratories Inc, Cat# PK6100 |  | 1:800 |
| **Visualization** | DAB–nickel substrate solution | 0.5mg/mL 3′,3′-diaminobenzidine-tetrahydrochloride, ammonium nickel sulfate, 0.01% hydrogen peroxide (H_2_O_2_) in TBS |  |  |

Notes: BBB, blood brain barrier, Cat#, catalogue number; IOD, Integrated optical density; MW, Microwave 800w for 10 min; NIN, Netherlands Institute for Neuroscience; TBS, Tris-Buffered Saline; PMID, PubMed unique identifier; RRID, Antibody registry ID
