## Supplementary figures and images for "Increased numbers of CD4^+^ T-cells in the hypocretin/orexin region of Narcolepsy Type 1"

### Supplementary Figure 1

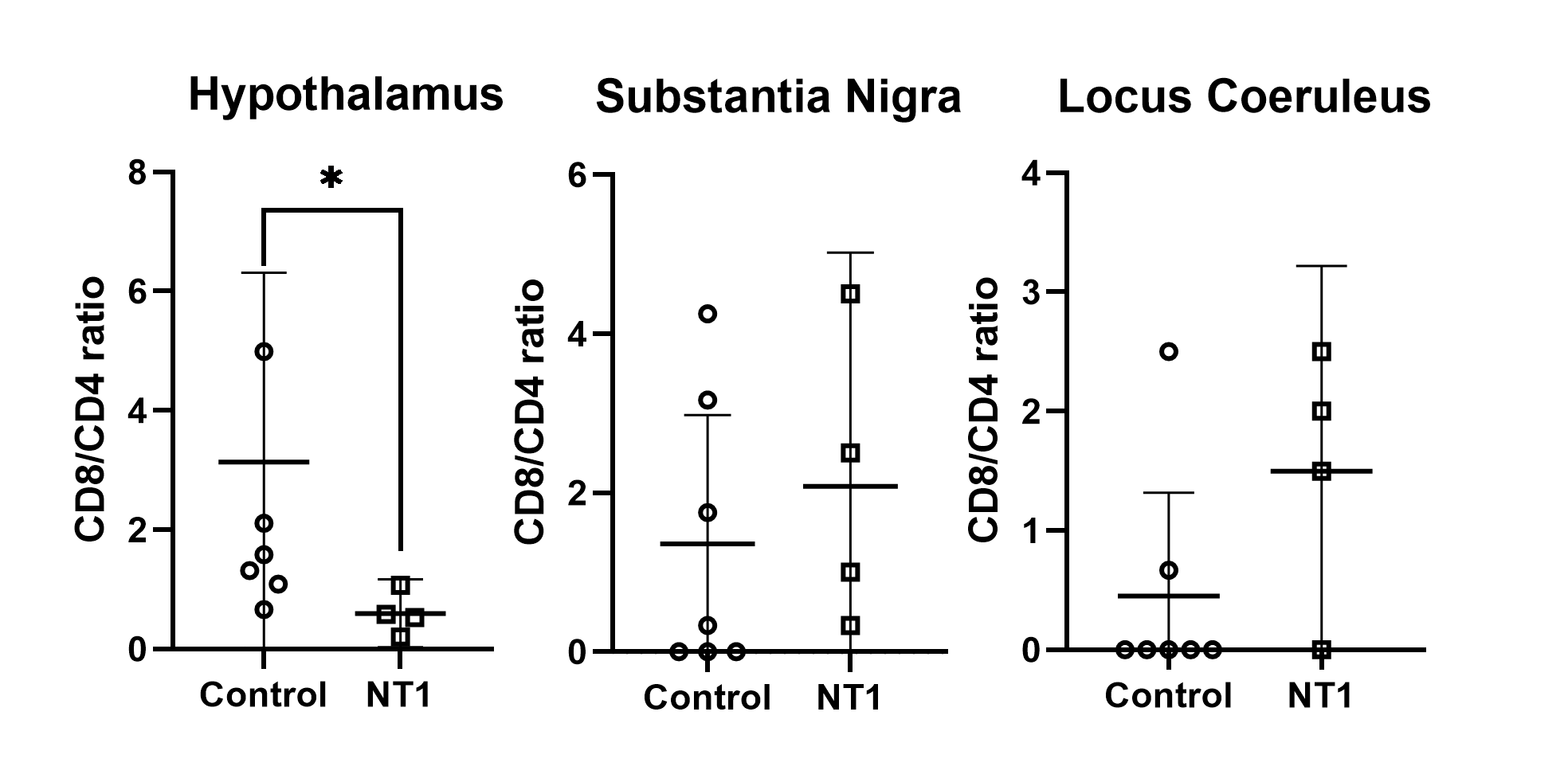
